# Tissue self-organization based on collective cell migration by contact activation of locomotion and chemotaxis

**DOI:** 10.1101/411306

**Authors:** Taihei Fujimori, Akihiko Nakajima, Nao Shimada, Satoshi Sawai

**Affiliations:** Graduate School of Arts and Sciences, University of Tokyo, Komaba, Meguro-ku, Tokyo 153-8902, Japan; Research Center for Complex Systems Biology, University of Tokyo, Komaba, Meguro-ku, Tokyo 153-8902, Japan

## Abstract

Despite their central role in multicellular organization, navigation rules that dictate cell rearrangement remain much to be elucidated. Contact between neighboring cells and diffusive attractant molecules are two of the major determinants of tissue-level patterning, however in most cases, molecular and developmental complexity hinders one from decoding the exact governing rules of individual cell movement. A primordial example of tissue patterning by cell rearrangement is found in the social amoeba *Dictyostelium discoideum* where the organizing center or the ‘tip’ self-organize as a result of sorting of differentiating prestalk and prespore cells. Due to its relatively simple and conditional multicellularity, the system provides a rare case where the process can be fully dissected into individual cell behavior. By employing microfluidics and microsphere-based manipulation of navigational cues at the single-cell level, here we uncovered a previously overlooked mode of *Dictyostelium* cell migration that is strictly directed by cell-cell contact. The cell-cell contact signal is mediated by E-set Ig-like domain containing heterophilic adhesion molecules TgrB1/TgrC1 that act in trans to induce plasma membrane recruitment of SCAR complex and formation of dendritic actin networks, and the resulting cell protrusion competes with those induced by chemoattractant cAMP. Furthermore, we demonstrate that both prestalk and prespore cells can protrude towards the contact signal as well as to chemotax towards cAMP, however when given both signals, prestalk cells orient towards the chemoattractant whereas prespore cells choose the contact signal. These data suggest a new model of cell sorting by competing juxtacrine and diffusive cues each with potential to drive its own mode of collective cell migration. The present findings not only resolve the long standing question of how cells sort in *Dictyostelium* but also cast light on the remarkable parallels in collective cell migration that evolved independently in metazoa and amoebozoa.

## Introduction

One of the fundamental processes that underlie tissue patterning is spatial rearrangement and repositioning of cells according to their cell-types [1-3]. *In vitro* studies have demonstrated wide occurrence of cell-type dependent segregation in the mixture of cells dissociated from different tissues [4-6]. Such cell segregation has traditionally been explained based on differences in cell-cell adhesion force and surface tension in analogy to phase separation e.g. of oil and water where membrane fluctuations would drive rearrangement of relative positions of cells so as to minimize total free energy. Quantitative measurements in conjunction with mathematical modeling have successfully provided qualitatively accurate predictions of *in vitro* sorting patterns [7,8]. While such view of cell segregation does seem to hold for *in vitro* systems, the extent of their contribution *in vivo* remains to be questioned. In many cases, such a stochastically driven process appear not to hold, as cells are migratory[9,10], and segregation occurs rapidly without being trapped in metastable states. In primitive streak of chick embryo and limb bud, directed migration is the primary driving force of morphogenesis[11,12]. In zebrafish gastrulation, internalization of mesendoderm cells require Rac dependent directed cell migration [9]. These examples point to importance of specific directional cues and migration in cell segregation, however the exact navigational rules at the single-cell level and their linkage to the resulting tissue patterns are still largely undeciphered.

In the social amoeba *Dictyostelium discoideum*, upwards of 100,000 cells aggregate by chemotaxis to self-generated waves of extracellular cAMP[13-17] to form a multicellular mound. In the mound, cells differentiate into either prespore or prestalk cells that initially appear at random positions before being segregated to form a distinct prestalk tip region[3,18,19] – an organizing center that sits on top of a prespore cell mass (Fig. 1A). During this process, cAMP waves cease (Fig. S1A and B), prespore cells migrate radially while prestalk cells exhibit a combination of radial and centripetal movement toward the apical region (Fig. 1B). Several lines of evidence suggest importance of chemotaxis to extracellular cAMP in cell segregation [20-22]. A gradient of extracellular cAMP formed by a glass needle in a mound can direct prestalk cell migration [22], and over-expression of cAMP-specific phosphodiesterase (PDE) suppresses tip formation [20]. On the other hand, heterophilic adhesion molecules TgrB1 and TgrC1 [23,24] are also essential for tip formation [25]. Knock-out mutant of TgrC1 exhibits motility defects [26] as well as loss of developmental gene expression[25,27]. Moreover, application of antibody against TgrC1 to regenerating mounds suppresses prestalk/prespore segregation [28]. TgrB1 and TgrC1 are also known for their polymorphism, which results in kin discriminatory segregation during aggregation [24,29,30]. These lines of evidence suggest requirement for extracellular cAMP and TgrB1/C1 for tip formation, however how they dictate the cell segregation process remains to be resolved [3,31].

**Figure 1.**
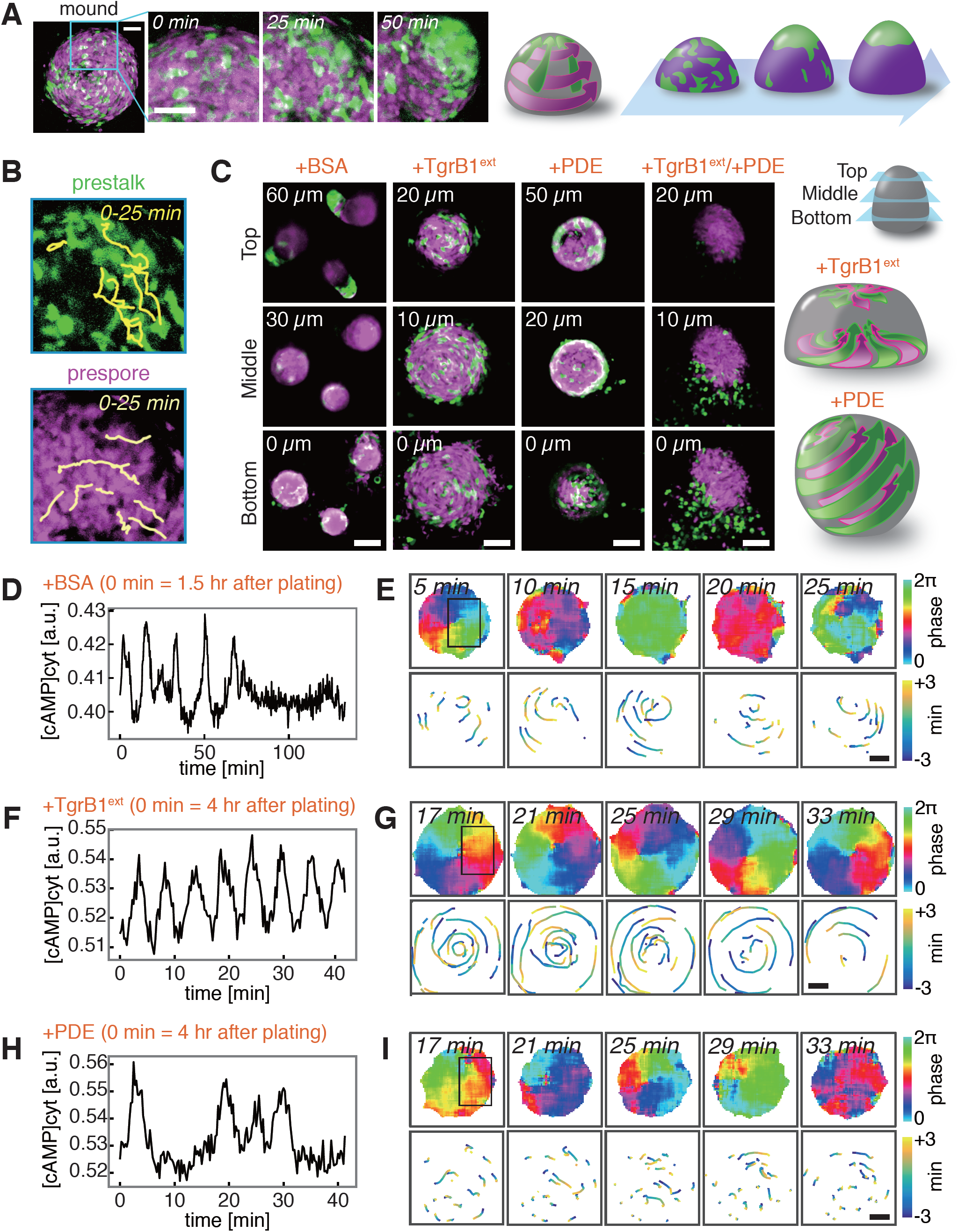
Two navigational cues and the modes of collective movement underlie segregation of prestalk and prespore cells in a *Dictyostelium* mound. **A**, Tip formation (*t* = 0, 25, 50 min. Green: prestalk marker ecmAOp:GFP, magenta: prespore marker D19p:RFP). Scale bars, 50 μm. **B**, Cell trajectories (upper panel, prestalk cells; lower panel, prespore). **C-I**, Interference of tip regenerating cues (Fig. S1E, Movie S1). Z-sections taken at 3 hr 40 min after plating (+BSA mock control, +TgrB1^ext^, +PDE, +TgrB1^ext^/+PDE)(**c**). Scale bars, 50 μm. Schematic illustrations of cell motion (right panel). **d-i,** Timeseries of the mean cytosolic cAMP levels (**D**, **F**, **H**) in boxed regions ((**E**), (**G**), (**H**) upper left panels). Cell trajectories (98% Epac1-camps/AX4[16]: 2% Lifeact-RFP/AX4 cells; +BSA (**D**, **E**), +TgrB1^ext^ (**F**, **G**), +PDE (**H**, **I**)). Phase of the cAMP oscillations (**E**, **G**, **I**; upper panels)[15]. Trajectories of RFP labeled cells during one cycle of the oscillation (**E**, **G**, **I**; lower panels). Scale bars, 20 μm.

## Results

### Navigational cues for *Dictyostelium* cell migration

To study how cell migration are being directed in the mound, we analyzed the effect of interfering with extracellular cAMP and TgrB1/C1. In order to circumvent developmental effects due to requirement of TgrB1/C1 on cell differentiation[25], we took advantage of the fact that the process is entirely self-organizing; i.e. it can be recapitulated by fully differentiated prestalk and prespore cells after dissociation[32]. Dissociated cells immediately began emitting cAMP waves, reaggregated and formed tips as cAMP waves ceased (Fig. 1C-E; Movie S1). When regenerating mounds were immersed in purified TgrB1^ext^ (Fig. S1C-F), cAMP wave propagation did not stop, and cells moved in highly-coordinated scrolling motion at least for the duration of our observation (Fig. 1F,G). Prestalk and prespore cells moved similarly and did not segregate (Fig. 1C, +TgrB1^ext^; Movie S1). When exposed to cAMP-specific phosphodiesterase (PDE) to attenuate extracellular cAMP, mounds became spherical, and the cells continued to migrate radially as the entire cell mass moved like a rolling ball (Fig. 1C, +PDE; Movie S1). Prestalk cells sorted out to the periphery but never collected to form the apical tip (Fig. 1C, +PDE; Fig. S1G). The rotational movement was not correlated with a few passages of residual waves (Fig. 1H,I) suggesting that cell migration, despite being highly coordinated, was not chemotactically oriented. When both purified TgrB1^ext^ and PDE were applied, prestalk cells were completely stalled while prespore cells retained some movement but were less coordinated (Fig. 1C, +TgrB1^ext^/+PDE; Movie S1). These observations indicate that, in addition to chemotaxis towards cAMP, there is an additional guidance cue mediated by cell-cell contact that directs collective cell movement.

To clarify the basic rule of cell movement, we analyzed migration of cells immediately prior to prestalk/prespore diversification (‘streaming-stage’ cells; see Methods) using a microfluidic gradient chamber (Fig. S2A,B). While moving towards the cAMP source, cells made head-to-tail contacts and formed trains (Fig. 2A; Movie S2). At low loading densities, most cell trains were short; many consisted of 2 cells (Fig. 2B). In both 2-cell and longer cell trains, leader cells formed lateral pseudopods and exerted exploratory trajectories similar to solitary migration, whereas those that followed were elongated, monopodal and moved ballistically (Fig. 2C; Fig. S2C-F, S3). To delineate the role of chemotaxis and cell-cell contact, response to a reorienting cAMP gradient was analyzed (Fig. S4). Within minutes after gradient reversal, solitary cells and leader cells changed their direction by extending *de novo* pseudopods (Fig. 2D, upper panels). While some follower cells also responded similarly, many exhibited no immediate response and continued to follow the cell in contact (Fig. 2D, lower panels; Fig. 2E; Movie S3). Given that the cells were treated here with adenylyl cyclase inhibitor SQ22536 [16,33]to suppress cAMP synthesis, these observations suggest that in addition to chemotaxis to cAMP required for the formation of cell streams[34], there may be an alternative mode of navigation that depends on cell-cell contact.

**Figure 2.**
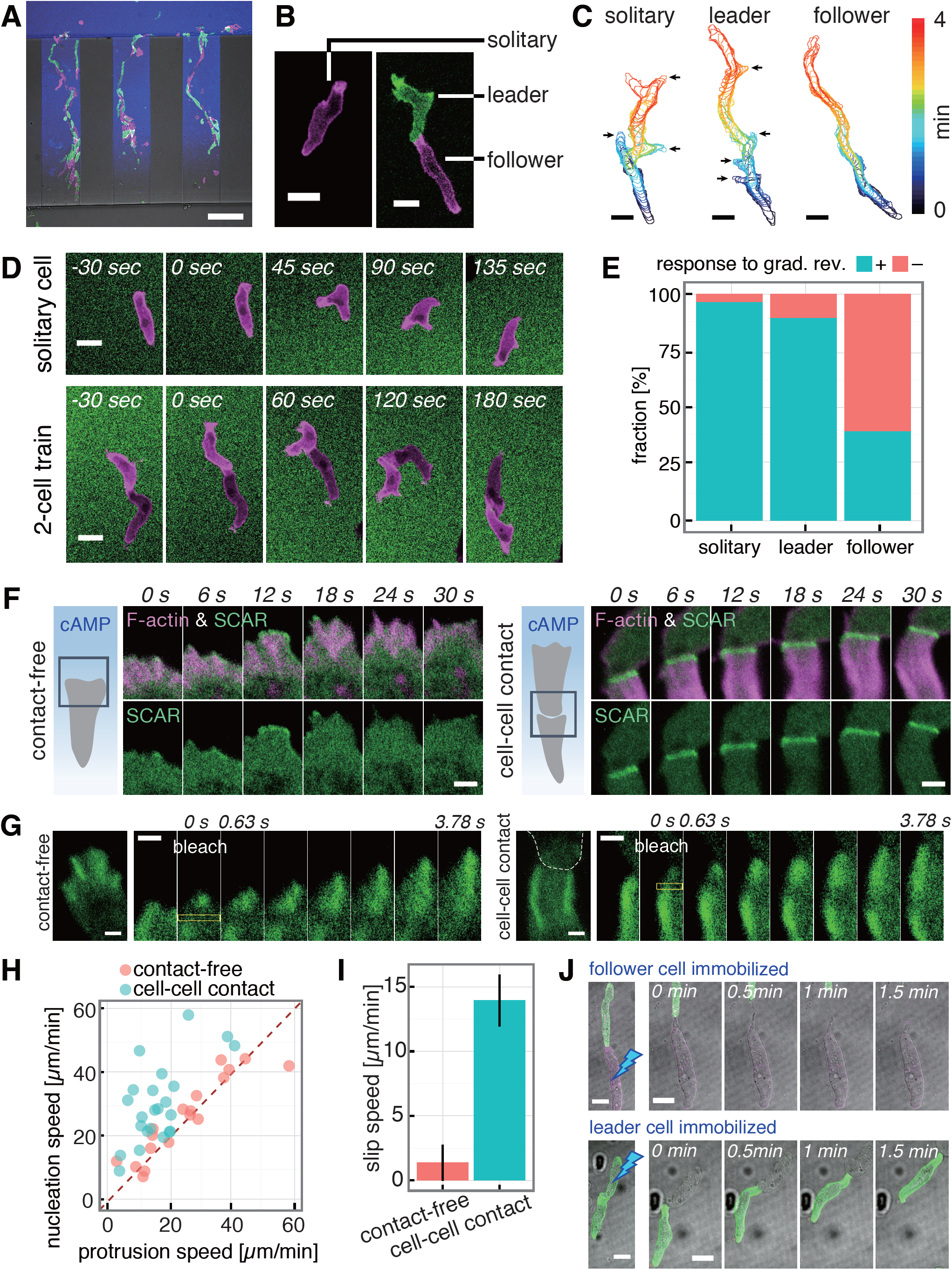
Microfluidics single-cell level analysis of train migration and contact-induced leading edge dynamics. **A**, Train migration. GFP-Lifeact/AX4 (green) and Lifeact-RFP/AX4 (magenta). 0-10 nM cAMP gradient (blue, ATTO425). Scale bar, 100 μm. **B**, Snapshots of solitary (left) and 2-cell train (right). Scale bar, 10 μm. **C**, Cell contours (arrows, lateral pseudopods). Scale bars, 10 μm. **D**, Response to reversal of cAMP gradient (0-1 μM; magenta, Lifeact-RFP; green, fluorescein). Scale bars, 10 μm. **E**, Fraction of cells with (‘+’) or without (‘–’) an immediate response (solitary: n = 73 cells, leader: n = 28 cells, follower: n = 97 cells). **F**, A contact-free (left panels) and a cell-cell contact (right panels) leading edge (green, HSPC300-GFP and magenta, Lifeact-RFP) in a 0-1 μM cAMP gradient. Scale bars, 2 μm. **G**, Fluorescence recovery after photobleaching in GFP-Arp2/AX4 cells (yellow boxed region) in a contact-free (left panels) and contacted (right panels) leading edge. White dashed line, a leading cell contour. 0-10 nM cAMP gradient. Scale bars, 1 μm. **H**, Scatter plot of nucleation and protrusion speed. **I**, Slip speed of actin filaments. Mean ± s.e.m., contact-free: n = 18 cells, cell-cell contact: n = 23 cells. **J**, Immobilization of a follower (upper panels) or a leader (lower panels) by UV irradiation. 0-10 nM cAMP gradient. Scale bars, 10 μm.

### F-actin dynamics at cell-cell contact sites

Compared to transient formation of F-actin at the leading edge without cell-contact (Fig. 2F, left panels, magenta)[35], F-actin at the cell-cell contact was persistent, long (∼5um) and appeared most strongly at the outer edge (Fig. 2F, right panels, magenta; Movie S4, Fig. S5A). The F-actin pattern was also observed in naturally streaming cells (Fig. S5B). The contact region was highly enriched in Arp2/3 (Fig. 2G) indicating that actin filaments form dendritic networks[36]. These observations are compatible with the general feature of a leading edge of migrating cells where dendritic F-actin networks grow mainly by side-branching nucleation mediated by Arp2/3 complex, while further away towards the cytosolic region, filaments are severed and depolymerized. The other widespread feature of the leading edge in various cells is the so-called ‘retrograde’ flow of F-actin due to excess filament growth relative to the speed of membrane expansion. Although solitary migrating *Dictyostelium* cells are known to lack obvious retrograde flow at the leading edge[37], timelapse images of F-actin at the cell-cell contact region were indicative of such flow (Movie S4). To quantitate the speed of retrograde flow of F-actin network, GFP-Arp2 incorporated in dendritic filaments was photobleached partially, and dislocation of the bleached region was followed over time. After photobleaching of GFP-Arp2, the non-fluorescent region moved backwards (Fig. 2G,H) at about 14 μm/min in cell-contacted leading edge compared to 1 μm/min (Fig. 2I) in cell-contact-free leading edge, suggesting an enhanced nucleation at the contact-site.

The enhanced F-actin formation at the cell-contacted leading edge suggests that there is upregulation of Arp2/3 activity. In solitary migrating *Dictyostelium* cells, the SCAR complex which is required for full activation of the Arp2/3 complex translocated to the membrane in small patches that last no longer than ∼10 sec [35](Fig. 2F, left, magenta). In contrast, at the cell-contacted leading edge, there was markedly enhanced localization of the SCAR complex (Fig. 2F; Movie S4). Major leading-edge signals such as Ras-GTP, PI(3,4,5)P3 and Rac-GTP were also present at the contacted-front, and appeared similar to contact-free leading edge (Fig. S5C-H). These observations suggest that the follower cells have a leading edge with persistent dendritic F-actin network that generates unidirectional propulsive force that pushes the plasma membrane forward. Accordingly, when leader cells were immobilized by UV irradiation, the follower cells continued to move and push the leader cells (Fig. 2J; Movie S5). The observation indicates contact-dependent protrusive activity that is independent of pulling by the front cell in contact.

### TgrB1/C1 and contact-induced front protrusion

The molecular basis of cell-train formation was further analyzed by studying binary mixtures of WT, *tgrB1*^−^ and *tgrC1*^−^. We found that in cells that follow *tgrC1*^−^, F-actin formation failed to become persistent (Fig. 3A,B; Fig. S6A,B). The pair-wise frequency of the contact itself was also low when cells being followed were *tgrC1*^−^ or when the following cells were *tgrB1*^−^ (Fig. 3C), which is consistent with a recent study[30] suggesting that TgrB1 and TgrC1 act as a receptor and a ligand, respectively, for allorecognition. TgrB1/C1 are developmental stage specific genes, hence the protrusions and the elongated shape in developing cells were never induced by contact with vegetative cells unless TgrC1 was overexpressed (Fig. 3D; Fig. S 6C,D). Moreover, streaming-stage cells were able to follow TgrC1 over-expressing vegetative cells that migrated towards their chemoattractant folate (Fig. 3E; Movie S6). Since vegetative cells do not secrete cAMP nor do the streaming-stage cells chemotax to folate, contact-dependent front protrusion and cell guidance are mediated primarily by TgrB1/C1 and does not require chemoattraction. Furthermore, silica beads coated with purified TgrC1^ext^ and lectin Wheat Germ Agglutinin (WGA) which binds to cell surface glycoproteins such as the homophilic adhesion protein CsA[38] were able to induce an extensive protrusion at the site of cell-microsphere contact (Fig. 3F). Much like cell trains, the cell-microsphere contact site was intensely decorated with the SCAR complex and F-actin (Fig. 3F; Movie S7). Microsphere coated with WGA alone or TgrC1^ext^ alone was unable to induce the characteristic protrusion (Fig. S6E). The results indicate that juxtacrine signaling between TgrC1 at the tail of a cell and TgrB1 at the front of a cell activates SCAR complex and induce highly enhanced formation of F-actin at the cell-cell contact site.

**Figure 3.**
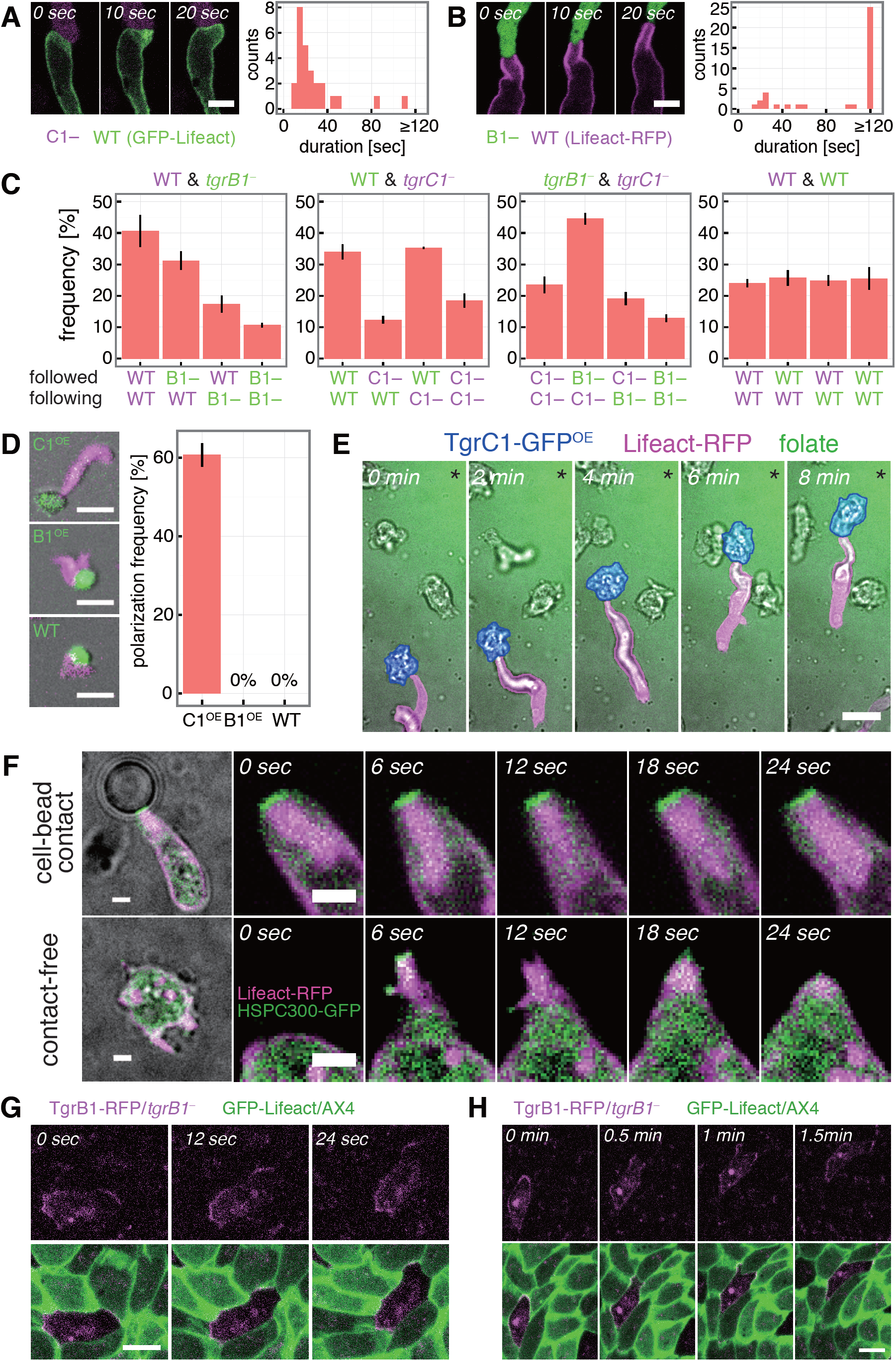
Contact-based front protrusion and train migration is mediated by TgrB1/TgrC1 interaction. **A-B,** Contact-induced F-actin formation (left; C1–: tdTomato/*tgrC1*^−^ (**A**), B1–: GFP/*tgrB1*^−^ (**B**), WT: AX4) and their duration (right; n = 26 events (**A**), n = 38 events **(B**)). Scale bar, 5 μm. **C,** Occurrence of head-to-tail contacts in 1:1 mixtures of GFP/*tgrB1*^−^ (B1–, green), tdTomato/*tgrC1*^−^ (C1–, magenta), Lifeact-RFP/AX4 (WT, magenta) and GFP-Lifeact/AX4 (green) (mean ± s.e.m.; n = 3 trials each; 120 ∼ 405 pairs total). **D**, Streaming-stage Lifeact-RFP/AX4 cells (magenta) attached to vegetative cells over-expressing TgrC1-GFP (upper left panel), TgrB1-RFP (middle left panel; CellTrackerGreen) or GFP-Lifeact (lower left panel). Occurrence of protrusions at the contact site (right panel; mean ± s.e.m.; n = 3 trials; total 49(C1^OE^), 46(B1^OE^), 16(WT) pairs). Scale bars, 20 μm. **E**, A streaming-stage Lifeact-RFP/AX4 cell (magenta) in contact with a vegetative-stage TgrC1-GFP over-expressing cell (blue) in a folate gradient (green, fluorescein; * source direction). Scale bar, 10 μm. **F**, Slug-stage HSPC300-GFP/Lifeact-RFP/AX4 cells attached to a TgrC1^ext^/WGA-coated microsphere (upper panels) or in isolation (lower panels) in a microchamber. Scale bars, 2 μm. **G-H**, A chimeric monolayer of TgrB1-RFP/*tgrB1*^−^ (magenta) and GFP-Lifeact/AX4 (green) in a microchamber. Scale bar, 10 μm. Splitting of a TgrB1-RFP enriched leading edge (**H**, 0.5 min) and selection of a single contact-site (**H**, 1.5 min).

Despite their spatially restricted mechanism of action, TgrB1 was localized to the front and the back of collectively moving cells whereas TgrC1 was observed uniformly at the plasma membrane (Fig. 3G; Fig. S7A-C). Lack of front/back symmetry breaking was puzzling considering that, in cell trains and aggregates, cells did not form two fronts in the opposing directions. In cells attached to 2 coated beads, cells indeed formed a front protrusion on one bead, and the other bead was attached to their tail except in cases where protrusions faced the same directions and hence cells were double-headed in shape (Fig. S7D). These results suggest that a protrusion is inhibited from forming at the tail once cells polarize. Interestingly, the choice of a bead to which protrusion formed sometimes swapped between the two (Fig. S7E) suggesting that contact-mediated polarity is dynamically maintained and possibly mechanosensitive. Similar front competitions were often observed in a monolayer aggregate where TgrB1 first accumulated towards two cells which then split as the cells deviated, and a contact with one cell was selected (Fig. 3H; Movie S8).

### Single-cell level response to navigational cues

It has been hypothesized that prestalk cells sort to the tip by migrating fast and winning the chemotaxis race against prespore cells[21,31]. When assayed at the single-cell level, however, prespore cells migrated faster than prestalk cells in a 0-1 μM cAMP linear gradient (Fig. 4A). Similar results were obtained from tracking well-isolated dissociated cells in the initial phase of reaggregation mitigated of contact signal by TgrB1^ext^ (Fig. 4B). In a 0-10 or 0-50 μM linear gradient, both cell types halted at the same location in the channel indicating a similar response range (Fig. S8). How then can cell-cell contacts facilitate cell segregation? When attached to TgrC1^ext^/WGA-coated microsphere (Fig. 4C), a marked protrusion was observed in about 1/3 of the cell-bead interface regardless of the cell-type (Fig. 4D). While the prespore cells exhibited a single protrusion at the contact site, many of the prestalk cells had auxiliary protrusions (Fig. 4C, arrows; Fig. 4E; Movie S9). When exposed to a cAMP gradient, the protrusion of prestalk cells at the bead-cell interface often disappeared and new pseudopods formed towards the cAMP source whereas prespore cells remained attached to the beads and retained the polarity (Fig. 4F,G; Movie S10). The results indicate that prestalk and prespore can be oriented by cAMP and TgrB1/C1, however when both signals are presented, there is a dominance as to which directs their leading edge. Prestalk cells prioritize response to cAMP and thus are chemotactically navigated, whereas prespore cells favor TgrB1/C1 and are navigated by cell-cell contact. Taken together with the results demonstrating requirements for two cues TgrC1 and cAMP for tip formation (Fig. 1C-H), the radial trajectories and the head-to-tail alignment of prespore cells is best explained by the Tgr-mediated navigation, whereas prestalk cells deviate from the contact-mediated collective migration and chemotax to extracellular cAMP (Fig. 4H).

**Figure 4.**
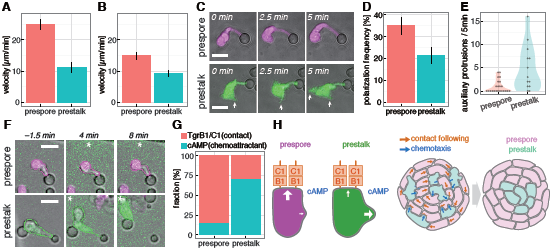
Single-cell level analysis of migratory response to navigational cues. **A**, Speed of isolated single-cells in 0-1 μM cAMP gradient (mean ± s.e.m., prespore: n = 33 cells, prestalk: n = 10 cells). **B**, Speed of single-cells during re-aggregation in the presence of purified TgrB1^ext^ (mean ± s.e.m., prespore: n = 28 cells, prestalk: n = 19 cells). **C**, A prestalk (green, GFP) and a prespore (magenta, RFP) cell attached to a TgrC1^ext^/WGA-coated microsphere. Prestalk cells form lateral protrusions (arrows). Scale bars, 10 μm. **D**, Frequency of cell-bead contact dependent polarization (mean ± s.e.m.; n = 3 trials). **E**, Number of lateral protrusion formed in cells attached to beads and elongated within 5 minutes (prespore: n = 23 cells, prestalk: n = 13 cells). **F**, Polarized cells attached to beads stimulated with a cAMP gradient (green, fluorescein; * source direction). Scale bars, 10 μm. **G**, Percentage of cells that maintain Tgr-mediated polarity (‘TgrB1/C1’) or abort the contact by protruding towards a cAMP gradient (‘cAMP’). Prespore: n = 19 cells, prestalk: n = 7 cells. **H**. A schematic illustration of the cell navigation rule.

## Discussion

A propensity of *Dictyostelium* cells to follow cells in contact was suggested by a classic work by Shaffer and coined the term ‘contact-following’[39], however it has remained heretofore unclear[40,41]. In the present work, we conclude that the following behavior is driven by ‘contact activation of locomotion’ – an induction of leading edge by cell-cell contact and the accompanying forward propulsion. The cell-contacted leading edge was highly enriched in a SCAR complex subunit HSPC300 and dendritic F-actin. Since the same response was observed in cells attached to TgrC1/WGA-coated microspheres, there appears to be a mechanism whereby TgrB1/C1 interaction induces accumulation of the SCAR complex at the cell-cell contact site. Lack of apparent features in the cytosolic residues in TgrB1 and requirement for lectin WGA for the response points to a possibility that the interaction between TgrB1 and the SCAR complex is indirect and that there is clustering of adhesion and signaling complex at the contact site. The contact site appears distinct from that forming a phagocytosis cup, since the contact area appears to extend narrowly instead of expanding and engulfing. Retrograde flow of F-actin has been suggested to play a major role in determining front-back polarity and its persistence during cell migration [42]. In addition, it has been shown that F-actin flow and persistent cell polarity is induced by confinement and decrease in cell-substrate adhesiveness [43]. The observations of the retrograde flow, monopodal morphology and loss of cell-substrate adhesion in the cell anterior are in line with these current understanding of the requisites for strong cell polarity in migrating cells.

Our results indicate that TgrB1/C1 has cell-type specific effect on cell polarity. It is essential for prestalk/prespore segregation as evidenced by well-mixed distribution of cells in a mound when contact signal was interfered with purified TgrB1^ext^ (Fig. 1C). While sorting of prestalk cells to the peripheral of the mound in the presence of PDE can be due to prestalk cells possibly being weakly cohesive [44], our data suggest that it accompanies their lesser ability to become monopodal and thus migrate directionally by contact. We should note that Myosin II accumulation at the plasma membrane has been shown to be stronger in prestalk cells than in prespore cells within a slug[45], and a null-mutant of myosin regulatory chain fails to form the mound tip [19]. Future studies are needed to clarify the molecular basis of cell polarity difference in prespore and prestalk cells.

The mechanism of collective migration in *Dictyostelium* uncovered in this study is in striking contrast to that of the neural crest cells. There, cell-cell contact signal mediated by Cadherin activates RhoA, inhibits protrusion and facilitate cell repulsion [46]. On the other hand, migration towards self-secreted chemoattractant C3a keep neural crest cells together [47,48]. In *Dictyostelium*, cell-cell contact mediated by TgrB1/C1 promotes protrusion, and chemotaxis rather is disruptive to otherwise more tightly packed cell mass as evidenced by mounds becoming spherical in the absence of the chemotactic cue. The present findings raise many open questions for future works. Besides prestalk segregation, the migratory mechanism may be relevant to *Dictyostelium* slug migration, culmination as well as kin-discriminatory segregation[24,29]. Also of note is a striking evolutionary convergence of collective cell migration, despite no homologues of TgrB1/C1 exist in metazoans. Are there parallelisms to Protocadherin-dependent SCAR complex recruitment and enhancement of migration in cultured cells[49] or similar enhancement of F-actin by atypical cadherin in rotating *Drosophila* egg chamber[50]? Are other cell streaming behaviors such as those observed in Human breast cancer cells[51] driven by a related mechanism? Further investigations in these phenomena should clarify common rules and logical necessities for cellular collectivity.

## METHODS

### Plasmid construction

His-tagged TgrB1 and TgrC1 expression vectors pA15-tgrB1^ext^-His_6_- 2H3term and pA15-tgrC1^ext^-His_6_-2H3term were constructed as follows. The expression cassette of pA15-GFP(S/T) was replaced with an expression cassette harboring the *act15* promoter, a *Cla*I/*Xho*I cloning cite and 2H3 terminator by *Xba*I and *Hin*dIII digest (pA15-2H3term). Complementary strands of synthetic His_6_-tag DNA oligos (Fasmac) were annealed and inserted into the pA15-2H3term vector at *Cla*I/*Xho*I sites by ligation. Genomic DNA fragments from the start codon to the codon just before the predicted transmembrane regions (*tgrB1*; Met1 to Thr802. *tgrC1*; Met1 to Asn851) were PCR amplified using Phusion polymerase (New England Biolab). The cloned fragments were verified by DNA sequencing and inserted into the *Cla*I cite of pA15-2H3term using Infusion HD cloning kit (Takara Z9633N) according to the manufacture’s protocol except that reaction volume was 5 μL containing 1μL premix, and the reaction time was extended to 1 hr.

TgrB1-RFP TgrC1-GFP expression vectors were constructed as follows. The *tgrB1* gene including the promoter and the ORF except the stop codon was amplified with an additional N-terminal *Nhe*I site and a C-terminal (G_4_S)_2_ linker sequence[52] and a *Cla*I site by PCR. Vector pHygTm(+) was cut by *Xba*I/*Cla*I to replace the *act15* promoter with the amplified *tgrB1* fragment to obtain pHyg-[tgrB1p]:tgrB1-(G_4_S)_2_-mRFP1. The vector with [tgrC1p]:tgrC1-(G_4_S)_2_-GFP was constructed similarly except that the *tgrC1* was cloned into pA15-GFP(S/T) for G418 resistance. TgrB1-RFP or TgrC1-GFP over-expression vectors were constructed by exchanging the native promoter in the plasmid with [tgrB1p]:tgrB1-(G_4_S)_2_-mRFP1 or [tgrC1p]:tgrC1-(G_4_S)_2_-GFP with the *act15* promoter. A prestalk GFP marker plasmid pEcmAO-GFP was constructed by replacing mRFPmars in pEcmAO-mRFPmars with GFP(S65T) at *Bgl*II and *Xho*I sites.

### Transformation and cell culture

Purified plasmids were used to transform *Dictyostelium* cells by electroporation following a standard protocol[53] and selected using 10 μg/mL G418 or 60 μg/mL Hygromycin. Following strains of *Dictyostelium discoideum* were used for live-cell imaging analysis: AX4 and its derivatives GFP-Lifeact[54]/AX4, Lifeact-mRFPmars[55]/AX4, GFP/*tgrB1*^−^, tdTomato/*tgrC1*^−^, GFP-Arp2/AX4, HSPC300-GFP[56]/Lifeact-mRFPmars/AX4, PakBCRIB-mRFP[56]/GFP-Lifeact/AX4, GFP-Lifeact/mRFPmars-Raf1RBD[17]/AX4, GFP-Lifeact/PHcrac-RFP/AX4, pEcmAO-GFP/pD19-RFP/AX4, Epac1-camps[57]/AX4, [tgrB1p]:tgrB1-RFP/*tgrB1*^−^, [tgrC1p]:tgrC1-GFP/*tgrC1*^−^, [act15p]:tgrB1-RFP/AX4, [act15p]:tgrC1-GFP/AX4. To obtain pEcmAO-GFP/pD19-RFP/AX4, AX4 cells were co-transformed by electroporation with plasmids pEcmAO-GFP and pD19-RFP. Phenotypes of TgrC1 or TgrB1 expressing clones in the respective knockout background varied on a bacteria plate likely due to stringency on the expression level. Clones whose null-phenotype of fruiting bodies were rescued were chosen for analysis.

Cells were grown in modified HL5 medium[15] with appropriate selection drugs at 22°C using a rotary shaker. To obtain the “streaming-stage” cells, cells were removed of nutrient by 3 repeated cycles of centrifugation at 700 ×*g* for 3 min and resuspension in Phosphate Buffer (PB: 12 mM KH_2_PO_4_, 8 mM Na_2_HPO_4_, pH 6.5). Washed cells were suspended at the final density of 5 × 10^6^ cells/mL and shaken for 6 to 8 hrs at 125 rpm at 22°C. cAMP pulses (final conc. 50 nM) were applied at 6 min intervals after 1 hr using a peristaltic pump (Masterflex L/S, Cole-Permer). To obtain “slug-stage” cells, washed cells were suspended in Developmental Buffer (DB: 6 mM KH_2_PO_4_, 4 mM Na_2_HPO_4_, 2 mM MgSO_4_, 0.2 mM CaCl_2_, pH 6.5) and plated on an agar dish (1% agar (Bacto) in DB) at final density of 6.6 × 10^3^ cells/mm^2^ to obtain a monolayer[16]. The cells were incubated in a dark condition at 22°C for 17 to 21 hrs before collection. Slugs were suspended in 1 mL of PB and mechanically dissociated by running it through a 23G needle (NN-2332R, Termo) back and forth using a syringe 10 times. Dissociated cells were washed and resuspended in 1mL of PB.

### Reagents

100 mM cAMP-Na (A6885, Sigma) solution was dissolved in water and stored at - 20°C as a master stock; working solution was 100 μM. Folate (060-01802, Wako) was dissolved in PB at 10 μM and used as a working solution. Fluorescein (231-00092, Wako) was dissolved in water at 1 mM and stored at 4°C. ATTO425 (AD425-21, ATTO TEC) and Alexa594 (A10438, Invitrogen) were dissolved in DMSO at 1 mg/mL and stored at –20°C. SQ22536 (505-44021, Wako) was dissolved in PB at 10 mM and stored at –20°C. Wheat germ agglutinin (WGA: 126-02811, Wako) was dissolved in water at 1 mg/mL and stored at –20°C. cAMP-specific phosphodiesterase (P0520-1UN, SIGMA) was dissolved in PB at 1 to 2 unit/mL and stored at 4°C.

### Microfabrication

Poly-dimethylsiloxane (PDMS) chambers were fabricated as described previously[58]. In brief, blank masks (CBL4006Du-AZP, clean surface technology, Japan) were UV-irradiated by laser drawing system (DDB-201-TW, Neoark, Japan) and chemically etched to create photomasks. Photoresists SU-8 3005 and 3050 (Microchem, USA) were used to create molds of the chambers. The thickness of SU-8 which determines the height of the microchamber for observation channel was approximately 4-5 μm for the gradient chamber (Fig. S2A and S4A), 4 μm for the low ceiling chamber for multicellular observations (Fig. S7B). The mask was placed on top of a SU-8 coated wafer and UV-irradiated using a mask aligner (MA-20, Mikasa, Japan). The cured SU-8 was used as a mold to fabricate PDMS. Glass cover slips (No. 1 or 1S, 24 × 60 mm or 50 × 60 mm, Matsunami) were washed in four steps using basic detergent, ethanol, NaOH solution, then rinsed in milli-Q water. Washed cover slips were dried at 140°C in a sterilization oven. The cover slips and fabricated PDMS were treated with air plasma, bonded and heated at 75°C for 1 hr.

### Flow chamber and stimulus delivery

Flow was controlled using a pressure regulator (Microfluidic Flow Control System (MFCS), Fluigent) or a syringe pump (70-4506, Harvard) as described previously[59]. A PDMS chamber was connected to the MFCS device with ethylene tetrafluoroethylene (ETFE) tubes. 5 μg/mL ATTO425, 3 μM fluorescein or 4 μg/mL Alexa594 was included in the cAMP solution to visualize the gradient profile. For assays of streaming-stage cells in the gradient chamber, 50 μM SQ22536[16,33] was included in the perfusion to inhibit adenylyl cyclase. Gradient was reversed by programmed flow control using a MFCS scripting module. For quantification in Figure 2E, timelapse series from 27 channels in total were acquired with 20× objective lens. Cells positioned at 95 to 378 μm from edge of the observation channel (0 to 473 μm) was analyzed. ‘Chemotaxis’ to gradient reversal was defined by formation of contact-free leading edge and migration toward a new gradient (Fig. 2E). Cell speed in Figures 4A and B was calculated from cell trajectories obtained by manually tracking cell centroid using ImageJ MTrackJ plugin. A glass needle (Femtotip, Eppendorf) was backloaded with either 100 μM cAMP or 10 μM folate solution in PB containing 10 μM fluorescein (Fig. 3E, Fig. 4F,G). To stimulate cells attached to a coated-microsphere, the needle was positioned approximately 20 μm from the cell of interest using a manipulator (TransferMan, Eppendorf), and cAMP gradient was formed by pressuring the needle at 80 hPa using an injection pump (Femtojet, Eppendorf)[60]. ‘Response’ to cAMP was defined by appearance of a protrusion toward the cAMP source and simultaneous loss of a contact-dependent protrusion (Fig. 4G).

### Microscopy and Image Analysis

All fluorescence images were obtained by confocal microscopy. With some exceptions (see below), data were obtained by an inverted microscope (Ti-E, Nikon) equipped with a laser confocal point-scanning unit (A1R, Nikon) using a Galvano scanner except for data in Fig. S5A where a fast resonant scanner was employed. For UV-irradiation, 405 nm laser (100 mW CUBE, Coherent, at max power) was employed. For photobleaching of GFP-Arp2 (Fig. 2F-H), 488 nm laser (100 mW CUBE, Coherent, at max power) was used to irradiate 0.25 × 4.88 μm (8 × 157 pixels) rectangular ROI positioned halfway between the ends of F-actin filaments. The length of actin filaments 1.26 sec and 3.15 sec after bleaching were measured manually, and the difference between them was defined as nucleation speed. The displacement of the tip of filament at the same time frames was defined as protrusion speed. Slip speed was calculated by subtracting protrusion speed from nucleation speed.

For data in Fig. 3E, 4F-G, an inverted microscope IX81 (Olympus) equipped with a multi-scan laser confocal unit CSU-X1 (Yokogawa) with EMCCD camera (Evolve512, Photometrics) was used. For data in Fig. 1C-I, 3D, 3F, and Fig. S1A-B, S6C-D, an inverted microscope IX83 (Olympus) equipped with CSU-W1 with a EMCCD camera (iXon 888, Andor) was used. 445 nm, 488 nm and 561 nm lasers were used for excitation, and fluorescence images were obtained by appropriate filters. Piezo z-stages were used for z-sectional imaging. Live-cell imaging was performed at 22°C. All images were stored as TIFF files and analyzed using homemade programs in ImageJ and MATLAB (Mathworks).

To obtain relative changes in the cytosolic cAMP level (Fig. 1D,F,H), the ratio of fluorescence intensities of Epac1-camps/AX4 in the CFP-channel (*I*_472/30_) and the YFP-channel (*I*_542/27_) was averaged in a rectangle region (24 × 36 pixels). For calculation of the phase by time-delay embedding (Fig. 1E,G,I), the mean fluorescence intensities in the CFP-channel and the YFP-channel images were averaged by moving timeframes (5 time points; 60 sec) and the background was subtracted. The background was obtained by averaging over 31 time points (450 sec) in moving time frames. The average fluorescence intensities 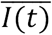cwere obtained for all positions by 5 × 5 pixels binning. Phase was defined by an angle in the embedding space 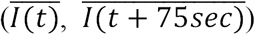. Lifeact-RFP expressing cells were manually tracked during ±3 minutes at each time points.

Quantification of cell morphology (Fig. S3), was performed as previously described[58] with some extensions. Velocity was obtained by calculating displacement projected normal to the cell contour per unit time and averaged over 3 neighboring positions and 3 time points. The sign of velocity was negative for inward projections. Localization of Lifeact-RFP or GFP-Lifeact at cell boundaries were calculated by taking the average fluorescence intensities for each 3 × 3 pixels regions at the boundary (rounded coordinates; 500 positions) and dividing it by the mean fluorescence intensities of the cytosolic region for normalization.

Duration of F-actin enrichment at the cell-cell interface (Fig. 3A,B, Fig. S6A,B) was measured based on the appearance of strong F-actin accumulation especially near the periphery of cell-cell contact. For the statistics of cell-cell contact, cells in contact for longer than 2 minutes were counted. For the analysis of head-to-tail pairing in binary cell mixtures (Fig. 3C), relative positioning of cell-types within cell trains were identified and counted manually based on the fluorescent labels. Frequency of pairing was corrected by total detected cell number of each mixed strain to eliminate the cell number bias. Cells attached to vegetative cells or a microsphere (Fig. 3D, Fig. 4D, Fig. S6E) was identified as polarized if the elongated shape at the cell-cell or cell-bead contact region lasted for more than 10 minutes. Protrusive structures from the lateral side of polarized cells that lasted at least 18 sec were counted as lateral pseudopods (Fig. 4E).

### Purification of TgrB1^ext^ and TgrC1^ext^ and microsphere coating

Cells harboring pA15-tgrB1^ext^- His_6_-2H3term or pA15-tgrC1^ext^-His_6_-2H3term were designed to secrete extracellular domain of TgrB1 and TgrC1, respectively (Fig. S1C). To verify secretion of recombinant proteins, TgrB1^ext^ and TgrC1^ext^ were extracted from growth medium by His Mag Sepharose excel (GE healthcare, 17-3712-21) according to the manufacturer’s protocol. The expected 150 kDa band was confirmed by SDS-PAGE and western blot using a 6xHis monoclonal antibody (MBL, D291-3) (Fig. S1D). For lab-scale production, the cells were grown shaken in modified HL5 medium for 2 days until they reached approximately 1 × 10^7^ cells/mL. The typical working volume was 660 mL split into two 1 L flasks. The medium was separated from the cells by centrifugation and passed through a syringe filter (0.22 μm or 0.45 μm). The His-tagged protein TgrB1^ext^-His_6_ or TgrC1^ext^-His_6_ was purified by affinity chromatography (AKTA pure 25M2, GE Healthcare) using a nickel-nitrilotriacetic acid (Ni^2+^-NTA) column (His Trap FF crude 5mL). Captured His-tagged protein was eluted in 20 mL of PBS with an optimized concentration of imidazole. 20 mL of the eluate was concentrated 10-fold using a spin column (Amicon Ultra 15mL 50K, Millipore), diluted in 15 mL of PB. The sample was further concentrated to 700 μL using the spin column by repeating the process twice. The final yield based on the absorbance at 280 nm was 0.1-0.6 mg/mL for TgrC1 and 1-1.5 mg/mL for TgrB1. Purified samples were stored in 1.5 mL tube at 4°C.

Purified TgrC1^ext^ was immobilized to functionalized silica beads (5 μm diameter, Sumitomo Bakelite BS-X9905) by covalent bond according to the manufacturer’s protocol. Briefly, 2 to 4 mg of dry beads were suspended in immobilization buffer and mixed with either 20 μg of the purified protein, 20 μg wheat germ agglutinin (WGA) or both. The mixture was incubated overnight at room temperature in a plastic tube gently rotated at 30 rpm. Beads were collected by centrifugation, and resuspended in 0.5 mL of PBS. The process was repeated two more times. The beads were then suspended in 400 μL of inactivation buffer in a plastic tube and gently agitated by rotation at 30 rpm for 1 hr. The inactivated beads were collected by centrifugation and suspended in 0.5 mL of PBS. Washing was repeated thrice, before finally suspending the beads in 0.2 mL of PB. The coated beads were stored at 4°C.

### Observations of regenerating mounds

To obtain a thin agar film, 200 μL of 1% Agar / DB was poured on a glass bottom dish, covered the glass (15 mm diameter) entirely, then removed to leave small residues typically of 30 μm height. 1 × 10^6^ mechanically dissociated cells were pelleted by centrifugation, then resuspended in 10 μL of PB or 1 unit/mL PDE or 1 mg/mL TgrB1 or 1% BSA. Cells were spread on a thin agar film using a pipette tip with care not to touch the agar, and allowed to settle for 10 minutes. A wet Kimwipe was included in the glass bottom dish, and the lid was closed during observation to avoid sample from drying.

## Data Availability

The data that support the findings of this study are available from the corresponding author upon request.

## Acknowledgments

The authors thank S. Hirose, A. Kuspa and G. Shaulsky for providing *tgrB1*^−^, *tgrC1*^−^, GFP/*tgrB1*^−^ and tdTomato/*tgrC1*^−^ strains and tgrB1/tgrC1 replacement vectors; M. Fukuzawa for pHygTm(+); R. H. Insall for PakBCRIB-RFP and HSPC300-GFP vectors; D. Knecht for GFP-Lifeact vector, V. O. Nikolaev and M. J. Lohse for Epac1-camps; Dicty Stock Center for pBig-GFP-Arp2 and pEcmAO-mRFPmars; Y. Shirokawa, M. Fujishiro and T. Sugita for technical assistance. This work was supported by MEXT KAKENHI Grant Numbers JP18H04759, JP16H01442, JSPS KAKENHI Grant Numbers JP17H0812, JP15KT0076, and in part by MEXT KAKENHI Grant Number JP17H05992, JSPS KAKENHI Grant Numbers 25710022 25103008 (to S.S.) and JSPS Grant Number JP16K18537 (to A.N.). T.F. was supported by a JSPS fellowship Grant Number JP17J08690.

## Author contributions

T.F., A.N. and S.S. performed prototyping experiments and conceived the study. T.F. planned and carried out plasmid construction, protein purification, cell transformation, image acquisition and analysis, wrote codes for image quantification. A.N. designed and constructed microfluidics chambers. N.S. contributed to protein purification. A.N. contributed to microscopy and microfluidics setup. S.S. oversaw and supervised all aspects of the project. T.F. and S.S. wrote the manuscript.

